# Predicting optimal growth temperatures of bacteria using learned structural information from a single protein

**DOI:** 10.64898/2026.06.15.732269

**Authors:** Michael Hoffert, Dru Myerscough, Nicholas B. Dragone, Matthew J. Gebert, Jonathan J. Silberg, Noah Fierer

**Affiliations:** Ecology and Evolutionary Biology, University of Colorado Boulder, Boulder, Colorado, United States of America; Cooperative Institute for Research in Environmental Sciences, University of Colorado Boulder, Boulder, Colorado, United States of America; Department of Bioengineering, University of California San Francisco, San Francisco, United States of America; BioSciences, Rice University, Houston, Texas, United States of America

## Abstract

Temperature is a fundamental determinant of bacterial physiology and ecology. Optimal growth temperature (OGT) is highly variable across species, contributing to differences in where and when species are most likely to thrive. Although the OGTs for most bacteria remain unknown, the increasing availability of genomes from uncultivated and cultivated taxa has made it advantageous to build genomic, cultivation-independent models to infer OGT. However, pre-existing genomic models often lack the generalizability and mechanistic grounding required for robust inferences of OGT. We propose a novel framework for predicting bacterial OGT which uses learned protein structural signatures of thermal adaptation. We hypothesize that biophysical tradeoffs which dictate enzymatic functions across variable temperatures provide a more robust empirical basis for OGT prediction than broad genomic features. Our OGT-predicting model, ROSEATE, is based on a single gene, adenylate kinase (ADK), that encodes for a ubiquitous enzyme essential for energy homeostasis. ROSEATE uses high-dimensional latent space encoding via MSA Transformer, a protein language model which embeds ADKs in a manner which preserves biophysical information about embedded proteins. We show that the accuracy of the ROSEATE model is on par with other genome-based models, has a high degree of phylogenetic generalizability, and the ESM embeddings effectively capture key temperature-adaptive enzyme characteristics derived from AlphaFold structures. Because ROSEATE is based on analyses of a single ubiquitous protein, it can be used with metagenomic data to infer the community-level variation in bacterial OGTs. We demonstrate this feature of ROSEATE by reconstructing ADK sequences from over 500 environmental and host-associated metagenomes, successfully distinguishing community-wide thermal preferences across diverse habitats, from polar oceans to mammalian guts. By transitioning from genomic proxies to informationally dense protein structural features, this work provides an efficient, interpretable tool for predicting bacterial OGTs across taxa and whole communities.

**Author Summary:** The temperature preferences of bacteria are key to determining where species are most likely to grow and how bacterial communities may respond to changes in temperature regimes. Unfortunately, the optimal growth temperatures of most bacteria, including a broad diversity of bacteria found in many host-associated and environmental systems, currently remain unknown as many bacterial species cannot be grown or studied in a laboratory. While we now have genomic data for many bacteria, using these data to infer optimal temperatures for bacterial growth has remained a persistent challenge.

We developed and validated a novel approach to predict bacterial temperature preferences. When heated, proteins often unfold, becoming nonfunctional. To adapt to warmer environments, organisms evolve more stable proteins which resist denaturing at high temperatures. Instead of analyzing a bacteria’s entire genome, our approach uses a protein language model to quantify stability-enhancing changes in a single protein found across all bacteria. We found that this single-protein approach can be used to effectively predict the optimal growth temperatures of individual bacterial species and even whole bacterial communities. By changing how we use genomic information to predict temperature preferences, our framework provides a scalable blueprint for predicting other important bacterial traits from protein structure information.

## Introduction

Temperature has a powerful and well-studied influence on ecological, physiological, and molecular dynamics in biological systems. This is particularly true for bacteria, whose activities and distributions are strongly affected by temperature [1–5]. Despite the abundance of studies quantifying the temperature sensitivities of individual taxa [6–8] and communities [9–12], our current knowledge of bacterial temperature preferences is derived almost entirely from the minority of bacteria which have been cultivated and have had optimal growth temperatures (OGT) measured via *in vitro* experimentation. Therefore, the temperature preferences for the broad diversity of bacterial taxa found in many host-associated and environmental systems that remain uncultivated [13] are unknown. Resolving this knowledge gap is essential to understanding how the structure, composition, and activity of bacterial communities change in response to shifts in temperature and ultimately predicting how temperature modulates the ability of individual bacterial taxa to grow, survive, and compete. [8,14–17]. Being able to infer the temperature at which bacterial growth is optimized is beneficial in many scenarios, from efforts to culture new species [18], discover biotechnologically relevant species [19], identify potential pathogens [20], and predict microbial responses to climate change.

Adaptations to temperature which could serve as predictors of OGT across both cultivated and uncultivated species are reasonably well-studied. By altering membrane fluidity [21,22], protein thermostability [23–27], and DNA and genomic structures [28–31], or via thermosensory, regulatory [32–34], and metabolic adaptations [35,36], organisms maintain proper function of biomolecules as temperature alters the molecular dynamics of cells. Higher temperatures and molecular kinetic energy can disrupt interactions between atoms that determine the structure, function, and stability of all biomolecules [37]. Thermostable proteins must maintain proper flexibility while avoiding denaturation, membrane lipids must be sufficiently fluid or rigid to allow passage of nutrients and sensory signals, DNA and RNA should not melt or be supercoiled, and metabolic reactions and intermediates must not react too slowly or quickly. Maintaining this delicate homeostatic balance requires many mechanisms which are detectable through experimental or observational means, allowing us to recognize and even discriminate between organisms with different temperature optima [24].

Despite the numerous mechanisms of temperature adaptation that could serve as indicators of OGT, the OGT of uncultivated taxa cannot be estimated using features which can only be measured *in vitro*. Given this limitation, most models which predict bacterial OGT use properties that can be measured or inferred from genomic data. These genomic models have the advantage that they can use rapidly expanding databases of bacterial genomes [38], including genomes assembled from metagenomes (MAGs), to infer OGT even for taxa that remain uncultivated. Genomic OGT-predicting models use broad genomic properties [39], the sequence composition of particular features, like tRNAs and rRNAs [40,41], and, most commonly, amino acid frequencies from proteomes [42–48] to detect increased frequency of components which contribute to thermostabilizing interactions, like ILYWVER residues and GC bonds in proteins and RNAs [24,49]. The presence of thermostabilizing interactions, particularly in protein structures, can be sensitive indicators of OGT because proteins require stable structures to function, but the stability of most proteins is marginal – that is, only minimally sufficient to maintain stability under a species’ selective conditions [50]. This “marginal stability” is caused by mutations occurring in high-dimensional protein sequence space, and prevents proteins from becoming excessively thermostable, instead producing continuous structural relationships with OGT that have been observed across functionally important proteins in many organisms [51–55].

Here we explore the use of explicitly quantified protein structural data for predicting bacterial OGT, aiming to increase the generalizability, interpretability, and data efficiency of predictive OGT models. Although biophysical properties which influence protein thermostability, like residue packing and surface areas, are assumed to be the causal factor behind OGT predictors like dipeptide frequencies [24,56], precise computation of these biophysical properties requires three-dimensional structures or, at the very least, structural information [57]. Additionally, the distinctive dipeptide frequencies of bacterial lineages with specific temperature preferences are often due to shared evolutionary history rather shared adaptation to temperature itself [58] and inclusion of these phylogenetically associated predictors causes models to suffer when applied to clades phylogenetically distant from training data. Finally, requiring complete genomes as input to a predictive model means that OGT inference for uncultivated taxa requires assembling high-quality genomes from metagenomes, a challenging task in many diverse bacterial communities.

We hypothesize that the use of predictors derived from recently developed structural biology tools can address the collective limitations of existing genomic models of OGT. More specifically, we hypothesize that by quantifying signals of temperature adaptation in single protein structures, we can create an efficient and biophysically interpretable method to estimate OGT. Here we focus on the structure of adenylate kinase (ADK), a small three-domain protein (Fig S1) which is a critical and highly conserved component of energy homeostasis in all cells [59] and a well-studied model of temperature adaptation [27,60–64]. Although ADK structures predicted using AlphaFold can capture features canonically associated with bacterial OGT (Table S1), the high residual variation in these structural characteristics make them insufficient for predicting OGT (Fig S1). Promising alternatives to predicting OGT from pre-determined structural characteristics are protein language model (PLM)-based approaches, as recent work has demonstrated that PLMs can learn structural information [65,66], and that latent representations, or embeddings, constructed by PLMs can be used to distinguish between psychrophilic and thermophilic proteins [67], including the catalytic parameters of ADK specifically [68,69]. For this study, we embed ADK sequences using MSA Transformer, a pretrained PLM [70], fitting site-specific regressors on these embeddings to predict OGT from ADK protein structural variation. We validate our new method, ROSEATE (**R**egression **O**n **S**ite-specific **E**mbeddings for **A**DK-based OG**T E**stimation) by comparing MSA Transformer embedding components to the known structural features we have computed from AlphaFold structures. ROSEATE improves the efficiency, interpretability, and generalizability of OGT-predicting models by using derived structural information for a single protein, critically improving the capacity of OGT models to be applied to poorly characterized taxa and incomplete genomes which are inevitably common in environments, including soils and sediments, which contain a high diversity of novel bacterial taxa [71].

## Results

### Development of PLM-based structural predictor

We developed ROSEATE using the Gosha database [72], an OGT dataset containing over 9,000 unique bacterial genomic records associated with OGT values. By matching each genome in Gosha to a representative from the Genome Taxonomy Database (GTDB) [73], OGT values were associated with phylogenetic information and genome sequences, from which we extracted the ADK variants. Partitions of this dataset, which contained ADK sequences paired with OGT values, were used in model training and testing (Fig S2A). Because the Gosha database is heavily biased towards mesophiles, with 95% of genomes having a reported OGT between 20 and 40 °C, and 67% having OGT of 28, 30, or 37°C (Fig S2B), evenly sampled subsets of this database were used in model training (Fig S2B, see Methods). We note that the Gosha records are not clearly curated to confirm “growth” versus “optima” and show discretization in 5°C increments when compared to growth curve derived OGT from [7], (Fig S2E), but this dataset is among the most comprehensive and curated datasets available for developing and testing genome-based models of temperature preferences.

To embed the ADK sequences using the multiple sequence aligned (MSA)-based ESM model MSA Transformer [70], all annotated ADK sequences were first aligned into a MSA using an outgroup of curated ADK sequences (see Methods). This alignment was provided to MSA Transformer for embedding. The attention framework in MSA Transformer uses information both within proteins (“row-wise”) and at aligned sites across proteins (“column-wise”) to compute embeddings which capture protein structural variance. Embeddings computed from layer 12 of the 12-layer model (esm_msa1b_t12_100M_UR50S) were used as inputs for multiple model architectures (Table S2). To test how data quality and quantity influence the models, each model was trained or tested using two mutually exclusive partitions of quality-controlled proteins from the OGT dataset (Fig S2), referred to as “C” for “Corkrey” (which only includes Gosha OGT values from bacteria with known, validated growth curves) and “E” for “Even” (which includes evenly subsampled OGT records from the entire Gosha dataset, see Methods). The Even/E partition contained 1,227 records, and the Corkrey/C partition contained 230. Generally, using the larger training dataset enhanced performance despite the potential inclusion of lower- quality observations. RMSE was between 8 and 11°C and R^2^ between 0.44 and 0.75 for all models, indicating that regressor architecture is less important than distinctions between proteins in the underlying embeddings, consistent with previous literature [74–76] (Table S2). We also performed phylogenetically structured cross-validation (CV) to test model generalizability (see “Analysis of phylogenetic influence on prediction accuracy”). After testing a number of architectures, we selected a per-site ElasticNetCV-based regression for further analyses because it achieved the best combination of RMSE, R^2^, and phylogenetic CV performance while allowing interpretation of the importance of both sites within the protein and embedding components within sites. (Table S2).

Our approach, which we have named ROSEATE (**R**egression **O**n **S**ite-specific **E**mbeddings for **A**DK-based OG**T E**stimation), uses regularized regressions for the 768 embedding features at each residue of the ADK protein alignment (Fig 1), allowing it to account for different structural trends in regions of the protein which are more important to thermostability, like the core, lid, and NMP-binding domains. Additionally, the architecture allows convenient insertion of new proteins into the alignment for prediction. ROSEATE generally has higher errors at high and low temperatures, a common trend in genome-based OGT models, influenced by both sparser data for non-mesophiles and the possibility that protein evolution in psychrophiles is difficult to quantify [44,77]. To compare ROSEATE to another genome-based approach that is widely used, we retrained and tested TOME [45] on the ‘Even’ dataset used as ROSEATE’s training data. TOME uses genome-wide pairwise dipeptide frequencies to predict OGT, representing an existing class of approaches which use genome-based heuristics to measure “structure-proximal” information [24,48,53,56]. Despite using at least one thousand times fewer residues per genome (215 residues per ADK in our model versus whole proteomes with a mean of 1.2 million residues), ROSEATE is nearly as accurate, achieving an R^2^ of 0.72 and RMSE of 9.7°C (Fig 2A) compared to TOME (R^2^ = 0.86, RMSE = 6.9°C, Fig 2B).

**Figure 1.**
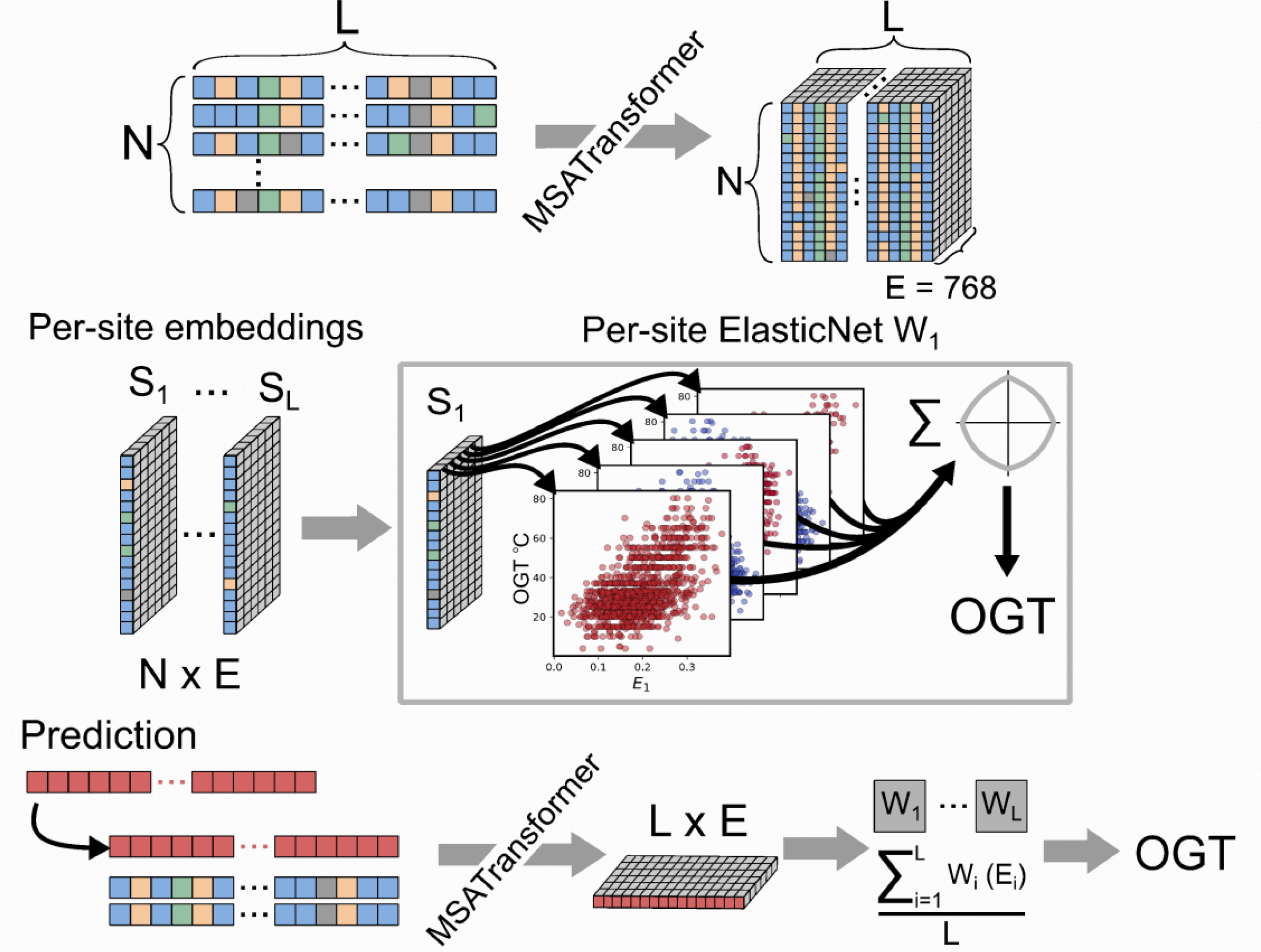
Overview of the ROSEATE model architecture. A N x L multiple sequence alignment is embedded using MSA Transformer [70]. Each of L sites across N proteins is embedded in a 768-dimension space. Each of these per-site embeddings is used to construct a per-site ElasticNet model which predicts OGT from embedded inputs. To predict OGT for a new protein, the protein is inserted into the original alignment, embedded, and embeddings from each site are passed to the site-specific models before averaging across sites.

**Figure 2.**
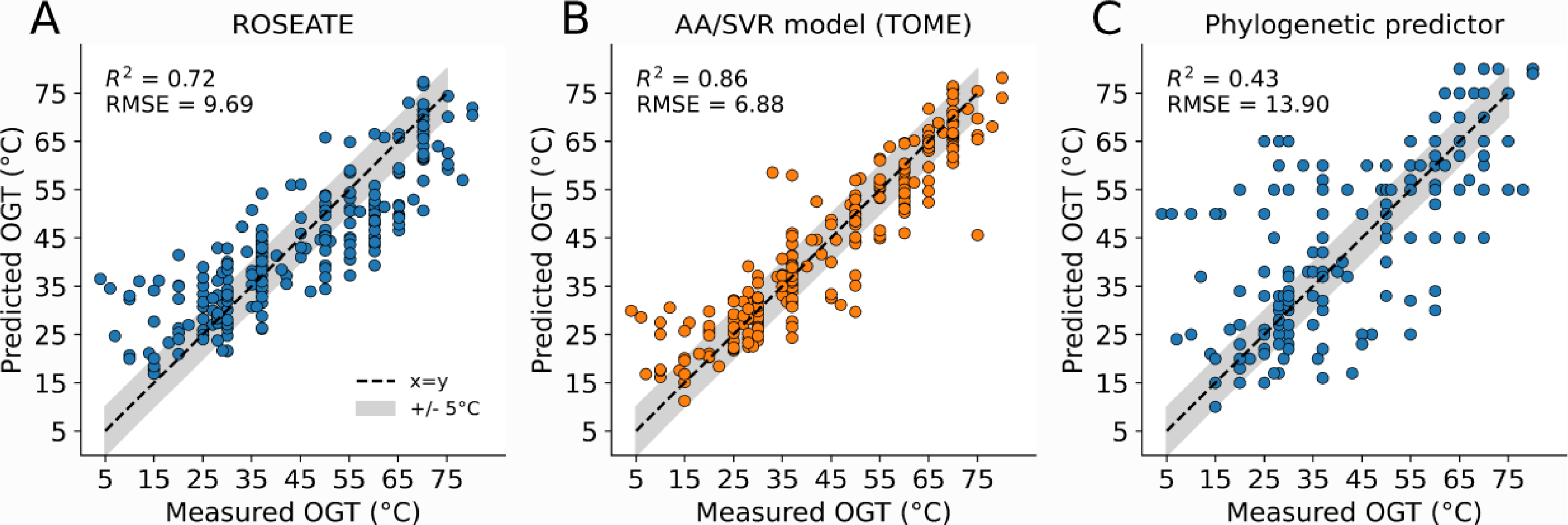
Performance of several model types for predicting OGT using genomic information. (A) ROSEATE, developed and described in this study, which stands for Regression On Site-specific Embeddings for ADK-based OGT Estimation, is an ESM-based model which uses 215-residue ADK variants to predict OGT. (B) TOME [45] uses support vector regression (SVR) and dipeptide frequencies calculated from whole genome sequences with on average 1.2 million residues included in the model. (C) A phylogenetic predictor, which predicts OGT for a species by simply predicting the OGT of its closest phylogenetic relative from a training dataset. All models use the same 1,227 training datapoints and 230 test datapoints (see Methods). Dotted line indicates a 1:1 relationship, with the grey band showing a 5 °C margin of error. R^2^ values and root mean squared error (RMSE) are indicated for each model.

### Characterization of novel isolate datasets

To assess the performance of these models on experimental data from closely related taxa, we applied both the ESM-based model and TOME to two new datasets of isolates not included in the Gosha database for which OGT values for each strain were independently determined by measuring growth responses across temperature gradients. The first isolate dataset contains twenty strains from the *Mycobacterium* genus [78] with measured OGTs ranging from 20-45°C. RMSE for ROSEATE and TOME differed by only 0.18°C, with ROSEATE achieving slightly higher R^2^ (0.21 vs 0.26, Fig S3A). The second novel isolate dataset contains 48 strains from the genus *Arthrobacter*, with 19 strains isolated from Antarctic soils and 29 isolated from soils outside the Antarctic continent, with measured OGT values ranging from 15-35°C [79]. Again, performance was similar (ROSEATE RMSE = 6.42 vs TOME RMSE = 5.98), with both models failing to distinguish a group of Antarctic isolates with average OGT of approximately 15°C from non-Antarctic isolates with OGT values closer to 30°C (Fig S3B).

### Analysis of phylogenetic influence on prediction accuracy

Machine learning methods like support vector regressions (SVR), used in TOME, cannot explicitly account for dependence structures introduced by phylogenetic covariance, which often lead to over-estimation of model performance, particularly when applied to taxa distantly related to those in the training data [80]. This is an important consideration given that the phylogenetic diversity of cultivated taxa is lower than in environmental and host-associated systems [81] where we want to apply ROSEATE. Our initial comparison of ROSEATE and TOME used a phylogenetic nearest-neighbors model (Fig 2C) to illustrate how OGT can be predicted for closely related taxa based purely on phylogenetic structure. Using the OGT of the nearest relative of a target taxon as its prediction, this phylogenetic predictor yielded a RMSE of 13.9 °C. Because closely related taxa (for example, members of the same genus) share evolutionary history, inclusion of taxa from the same genus in training and testing datasets effectively results in ‘data leakage’, where the training and testing data are non-independent. As a result, a model may ‘memorize’ features specific to a lineage rather than learning broader biological mechanisms, inflating its accuracy. Models can be evaluated for phylogenetically biased accuracies by assessing performance loss when taxa in training and test data contain a maximum degree of phylogenetic similarity [80].

To further examine the performance of ROSEATE and TOME when applied to taxa that are more phylogenetically distant from taxa included in the training dataset, we performed phylogenetic cross-validation. We retrained ROSEATE and TOME using a blocked phylogenetic cross-validation (PCV) approach applied to the ‘Even’ dataset (see Methods) to test for phylogenetic biases. The average PCV error of ROSEATE when training and testing datasets did not contain the same phyla was approximately 9.77°C (Fig S4A), similar to our unblocked test accuracy (9.66°C). ROSEATE’s PCV error at the genus level (∼6°C) was similar to error in our tests on genus-level groups of isolates (6.42°C), indicating that our tests accurately approximate ROSEATE’s accuracy even when it is applied to phylogenetic groups which are phylogenetically divergent from those in the training data. The selection of the ElasticNet for ROSEATE was influenced by this analysis, since the regularized ElasticNet had the lowest PCV error across phylogenetic levels compared to other model architectures. Although the PCV error of TOME is very low when blocked below the phylum level (75% of repetitions with RMSE <2.5°C, Fig S4B), the phylum-level PCV error was similar to our tests. Unlike ROSEATE, the estimated errors of TOME are inconsistent between the phylogenetic PCV procedure (Fig S4) and our testing (Fig 2), indicating that the dipeptide frequencies used in TOME may be more reliant on phylogenetic information to accurately predict OGT values. Such TOME being less useful when attempting to predict OGT for bacteria that are more distantly related to those taxa amenable to cultivation-based study for which pre-existing OGT data are available.

### Assessment of ESM embeddings using a structural dataset

An initial assessment of how ADK structural variants in our OGT dataset agree with known temperature adaptations was conducted by folding annotated ADK sequences with ColabFold v1.5.5 [82] and measuring 27 structural features drawn from literature sources on protein adaptation to temperature (Table S1). Because the residual variation of these features was very high (Fig S1), our approach to predicting OGT instead uses protein embeddings generated by MSA Transformer as inputs, which we still describe as “structure-based” because these embeddings stratify proteins according to various structural characteristics which cannot be inferred from sequence alone [65,66], presumably including those which contribute to thermal adaptation. Although PLM embeddings have been used to predict both protein thermostability and thermotolerance of organisms [77,83], few studies further compare PLM embedding spaces with measured, known features of thermal adaptation to determine whether these features drive the predictive performance of PLM-based methods and not sequence homology, compositional biases, or noise in the embeddings. To preclude this possibility in ROSEATE, we compared OGTs with MSA Transformer likelihoods and an identical “one-hot” model which substituted ESM embeddings for one-hot encodings at each protein site. Both of these approaches were weakly correlated with OGT (Fig S5), indicating that ROSEATE is not regressing based on residue identity or similarity to MSA Transformer’s training data. Furthermore, the accuracy of ROSEATE continuously increases as the embeddings pass through MSA Transformer (Fig S6), which mirrors previous observations that secondary structure and tertiary contact prediction improve in later layers of PLMs [65,84]. After observing that the performance of site-specific models in ROSEATE is clearly structured by the domain layout and secondary structures of ADK (Fig 3), we sought to specifically probe how ESM embeddings represent structural features which contribute to thermal adaptation in bacterial ADKs.

**Figure 3.**
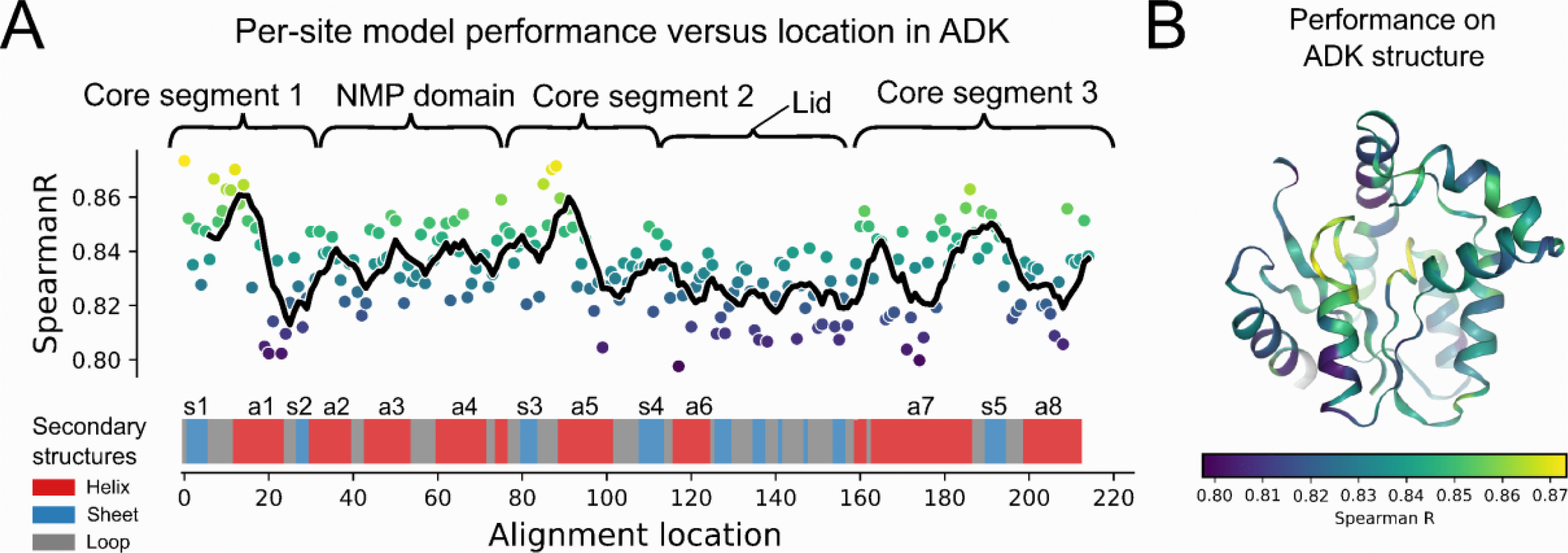
Performance of per-site models in ROSEATE. (A) Spearman correlation coefficients for each per-site model’s predictions in ROSEATE illustrating how particular sites in the protein, particularly those concentrated in the core domain, are more useful for predicting OGT. Alignment locations on the x-axis are calculated from backbone alignment of all ADK variants used for inputs to MSA Transformer. The secondary structures where each residue in the alignment resides are indicated above the alignment location. (B) AlphaFold-based ADK structure from *Dichotomicrobium thermohalophilum* (GTDB; RS_GCF_003550175.1) with lid excluded, colored by Spearman R values from (A) to illustrate the association of predictive accuracy with proximity to the active site and helix-sheet junctions in the core domain.

There is a notable concentration of highly predictive sites in two core domain sheet-helix junctions near the active site in the core domain generally, and within alpha helices in the NMP domain of the protein (Fig 3). This finding is consistent with prior biophysical studies showing that engineered ADK chimeras inheriting the core domain from a thermophilic ADK have higher thermostability than chimeras inheriting that domain from a mesophilic ADK [60], and lower degrees of tolerance for conformational flexibility within the core domain [27]. To identify which structural features have the strongest relationship with OGT in our dataset, we used a regularized linear model on 8,352 ADK structures retained after quality filtering to quantify the relationships between these structural features and OGT (Fig S7A). The relationships between structural features and OGT are largely consistent with previous findings (Fig S7B), with thermostability characterized by the presence of a zinc-binding lid domain [85], increases in solvent accessible surface area [86], and increases in contact density [87]. The major negatively correlated component was the score of the Rosetta energy function [88], an approximation of free energy of the native state which decreased in proteins from more thermophilic organisms as expected. The residuals of this regression below 20°C indicate that, consistent with ROSEATE, the measured structural features fail to resolve differences between proteins from putatively psychrophilic taxa (Fig S7C). Although studies frequently conclude that the underrepresentation of psychrophilic taxa is to blame for poor performance of OGT models below 20°C [47], some ADKs from putatively psychrophilic taxa in our dataset have substantially lower Rosetta reference energies (Fig S1D) and may be more thermostable, or from organisms with OGT values that are actually higher than reported in Gosha. We used the most highly associated structural features to further explore whether ESM embeddings capture these features.

At the protein scale, latent dimensions within the ESM embedding space are consistently aligned with established thermostability-associated structural features. The CLS token, which contains a protein-wide summary embedding, was correlated with average SASA, average number of contacts at 4.5Å, total Rosetta energy [88], and average ΔSASA, all protein structural features known to be associated with thermal adaptation [53]. With respect to OGT, each of these four structural features had directionally consistent associations with multiple embedding components (Fig 4). Embedding components positively correlated with features like SASA were also assigned positive coefficients in ROSEATE, meaning they positively predict OGT (Fig 4 A,B,D). The converse was true of negatively associated components and negatively associated features (Fig 4C). This alignment between the learned representations produced by MSA Transformer and explicitly characterized biophysical properties suggests that ESM-based models can innately stratify the protein landscape according to known mechanisms of thermal adaptation.

**Figure 4.**
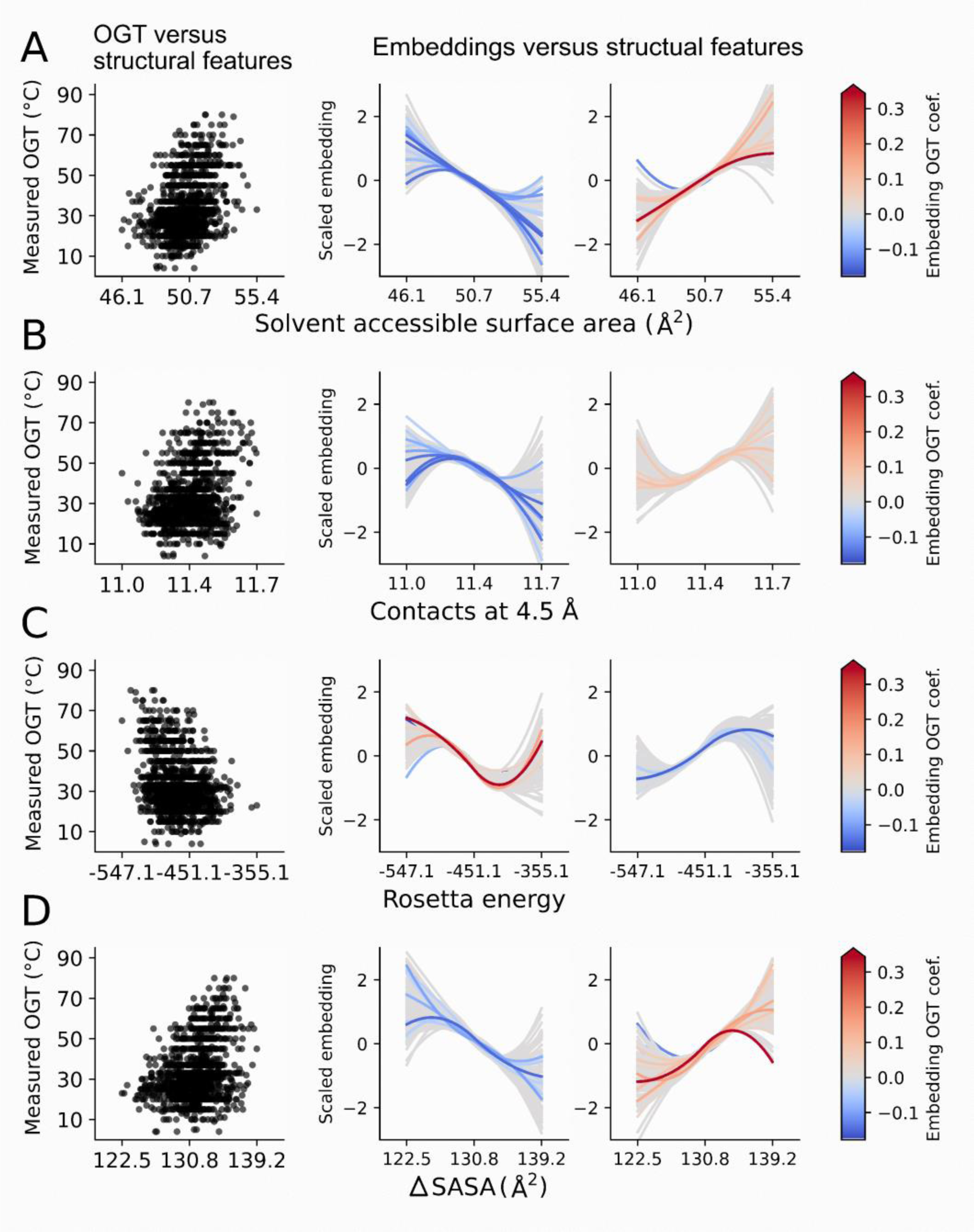
Associations of four key ADK structural features with OGT and MSA Transformer embedding components. First column shows values of each structural feature versus OGT. Second and third columns show trends in the relationship between embedding components (y axes) and each structural feature (x axes). For each structural feature (A-D), the 100 most negatively and positively correlated embedding components are show in columns two and three respectively. Each embedding versus structure trendline is colored by its coefficient in the ROSEATE model to illustrate how specific embedding features are correlated with structural features and derive some of their predictive power from their capacity to estimate biophysical characteristics of proteins known to be associated with thermal adaptation.

### Community-level OGT determination using metagenomic data

Rather than requiring complete assembly of genomes from metagenomic data, ROSEATE uses information from a single protein (ADK) to infer bacterial OGT. The distribution of OGTs inferred from the variants of ADK present in a sample reflects both the specific temperature preferences of organisms in that sample and the distribution of OGT values in the community. To test the application of ROSEATE on metagenomic datasets, samples from systems that we expect to vary in thermal regimes were collated and ADKs were extracted from each of the 415 metagenomes. The datasets included metagenomes from Australian soils [89], Antarctic soils [90], European freshwater lakes [91], cattle rumen [92], avian gut [93], hot springs [94], geothermally heated seafloor sediments [95], and ocean waters from polar [96] and nonpolar [97] regions. The distribution of estimated community-level OGTs across the eight environments broadly matches expectations (Fig 5). For example, the mean OGT values of bacterial communities from colder aquatic environments, both marine and freshwater, are lower than those from cattle rumen, geothermal sediments, avian guts, and hot springs, with the patterns closely aligning with environmental temperatures (Fig 5B). Likewise, the range in estimated OGTs within individual communities is higher in environments where we expect more variation in temperature regimes (e.g. soil) than in environments where temperatures are more stable over time (e.g. avian gut, Arctic Ocean) (Fig 5A). Together these results highlight how the single-gene architecture of ROSEATE enables efficient estimation of both community-level average OGT values as well as the estimated range in bacterial OGT values in individual samples. Future work could improve this approach via targeted assembly or amplification of ADK variants to enable OGT analyses of even rare community members from metagenomes while limiting the computational burdens of genome assembly.

**Figure 5.**
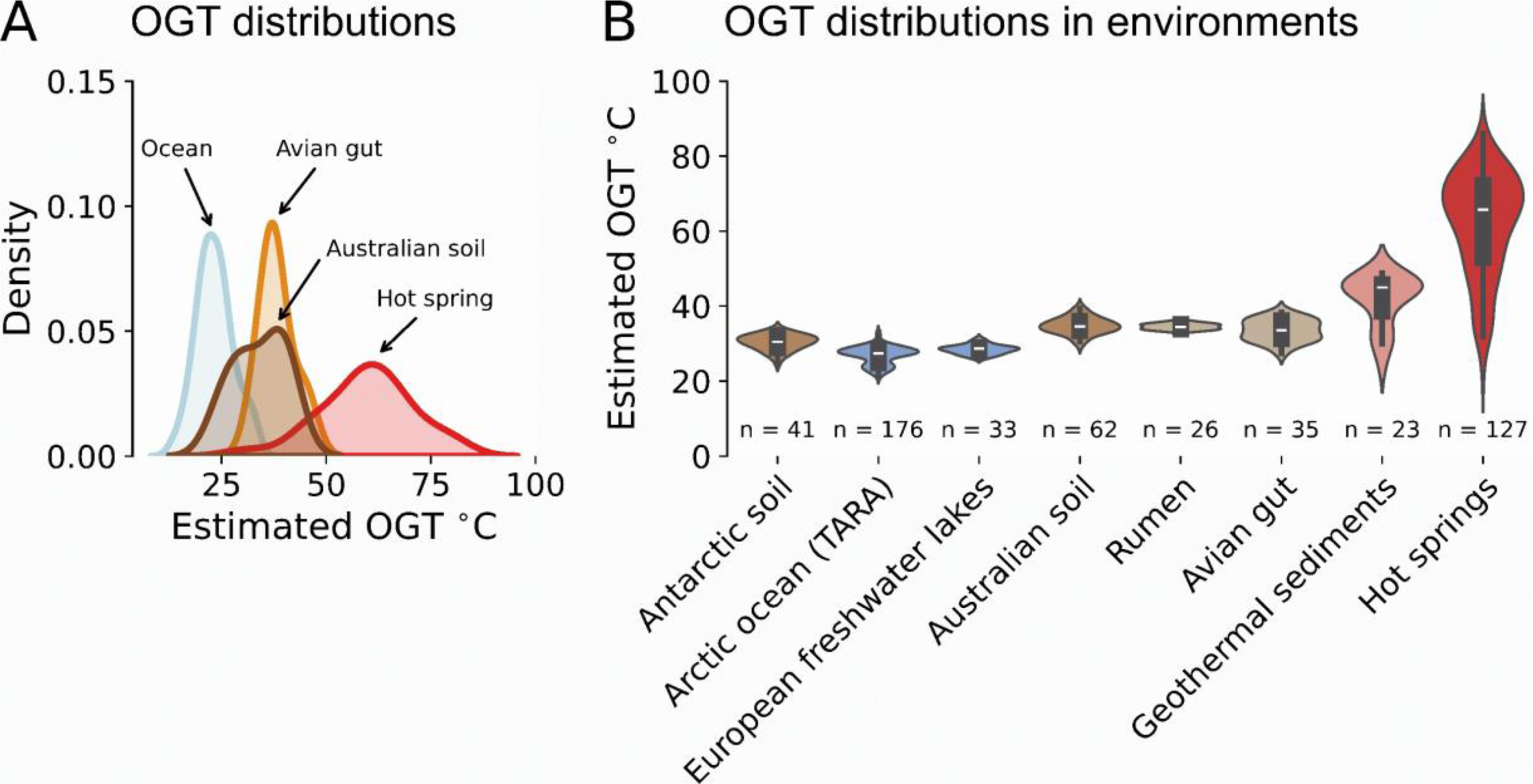
Application of ROSEATE to metagenomic data to predict the range and average OGT values of bacterial communities. (A-B) ADK sequences were assembled and annotated from shotgun metagenomes collected from samples with distinct environmental temperatures. (A) The ROSEATE model infers a distribution of OGT values in each sample by predicting the OGT for each ADK variant found in each sample. Figure shows results from four individual samples (metagenomes): an Arctic ocean sample (light blue), an avian microbiome sample (dark red), a hot spring sample (red), and an Australian soil sample (tan). (B) Distributions of estimated mean OGT values per sample for each of nine metagenomic datasets. The number of individual samples (metagenomes) included in each dataset is indicated for each type of environment.

Although estimations of OGT from our model generally match expected differences in temperature regimes (Fig 5), application of our model to these metagenomes is not a validation *per se* because the habitat temperatures are not necessarily related to the OGT of community members. Most notably, ROSEATE predictions for the Antarctic soil metagenomes are between 20 and 40°C, despite mean annual temperatures of -40 to -20°C for the sampling sites [79,90]. One explanation is simply that ROSEATE may not as accurately predict lower OGTs given the paucity of psychrophiles in the training dataset [47]. Alternatively, the unexpectedly high OGT values for bacteria found in Antarctic soils may be real, as the mean community-level OGT values (Fig 5B) are similar to the experimentally determined OGT values, ranging from 15 to 30°C, for members of the *Arthrobacter* genus isolated from these Antarctic soils (Fig S3B), results that are similar to those reported in other studies of Antarctic bacteria [79]. Bacteria living in cold environments may have appreciably higher OGT values than mean environmental temperatures as a bet-hedging strategy: while cold conditions are tolerable through dormancy, overheating is frequently catastrophic [98]. Another possibility is that bacteria in Antarctic soils may only grow during relatively narrow windows of time when soil temperatures are elevated. Although this study was not designed to explicitly test hypotheses about the relationship between bacterial OGT and environmental temperature, our results indicate that the range of OGT values within individual samples may be related to temperature variability, in addition to the mean differences in community-level OGT values capturing broader variation in temperature regimes across major environment types.

## Discussion

ROSEATE demonstrates that structural information can be efficiently incorporated into models which predict bacterial OGT through PLM embeddings, replacing sequence-level heuristics with embeddings which approximate structural information. Our work agrees with recent conclusions that ESM embeddings can be used for higher-order inference of protein properties [69] and convergent evolution [99], including of OGT specifically [77,100,101]. The primary limitations of our work and other PLM-based approaches are in the broad and unresolved challenge of interpreting PLM latent spaces. Few studies which use ESM to predict OGT attempt to define whether ESM embeddings reflect canonical biophysical patterns of thermal adaptation. While we have attempted to draw some comparisons between structures and embeddings as a starting point for such work, additional studies to validate whether the shifts in ESM embeddings reflect changes in the thermostability of proteins specifically and thermal fitness of organisms more generally are needed. Further knowledge of how PLM-based models identify causal influences on thermal fitness not only benefit efforts to validate OGT models but are also broadly relevant to efforts to infer microbial traits from genomic and metagenomic data. Without a means to experimentally validate predicted traits in most microorganisms, we believe that structural approaches like ROSEATE provide a novel and untapped source of empirical information about environmental preferences and the biophysical constraints on environmental tolerances.

The availability of high-quality OGT data for model training and validation is another persistent limitation shared by ROSEATE and other genomic models of OGT. The curated OGT dataset used here [72] and those used in other modeling efforts [48,102–104] are unavoidably biased towards bacterial taxa that are more readily cultivated and studied *in vitro*, with just four phyla (*Pseudomonadota*, *Actinomycetota*, *Bacteroidota*, and *Bacillota*) representing 90% of entries in our OGT dataset. Because OGT is frequently inferred from observations at specific temperatures rather than formal growth rate analyses across thermal gradients, OGT datasets lack the balance and resolution required for high-accuracy modeling. Many collections of OGT records are heavily discretized and disproportionately concentrated at 28, 30, or 37°C (Fig S2B), a distribution which is often blamed for underperformance of OGT models at high and low temperatures. In our work, data quality effects our determination of whether ROSEATE’s observed error of approximately 10°C represents an irreducible error stemming from data quality issues, a need to further refine the model architecture, or a true deviation of the temperature optima of proteins from our expectations. These limitations on phenotypic data for microorganisms create a bottleneck in predictive efforts like ours, which may not be primarily restricted by algorithmic capability but instead by the availability of high-quality training data. Therefore, while improving OGT model designs is important, expanding and curating OGT and microbial trait datasets is an even more critical step to ultimately improving machine-learning based predictive tools like ROSEATE.

Despite these limitations, we believe that structure-based OGT approaches like ROSEATE show significant promise for both clarifying methodological opacities and further improving efficiency, accuracy, and usability of OGT models. Further incorporation of structural biology methods could explore whether loss of predictive accuracy at low temperatures is due to thermodynamic effects, rather than data or model limitations. For example, low thermodynamic thresholds for folding at low temperatures may result in diverse sequences remaining folded, whereas the compromised hydrophobic effect at high temperatures may reduce viable strategies for thermostability to a small number of conserved, recognizable dynamics [105], making thermophily fundamentally easier to predict. In addition to insights into the biophysical dynamics of selection for thermostability, the use of protein structure and protein language models in our work also enables many future improvements, including the ability to expand to additional proteins, consider other stability-enhancing biophysical dynamics [106], and identify complex functional convergence in proteins [99]. These benefits merit comparison to recent OGT models, which only marginally develop on the use of dipeptide frequencies as OGT predictors from 2019 [45–48]. By substituting large volumes of weak genome-wide heuristics with concentrated representations of a single protein, we have found a way to improve how those inputs represent the sequence characteristics which predict OGT while reducing their size (and the associated work to construct them) a thousand-fold.

In summary, while ROSEATE’s accuracy might be limited in part by the quality and size of OGT datasets and the complexity of interpreting PLMs, it achieves comparable accuracy to existing methods while offering distinct advantages over pre-existing genome-based OGT models. These improvements come from the use of learned protein representations, which take advantage of a pretrained ESM model to reduce the number of sequences which must be reconstructed from metagenomic data. By focusing on a single protein, ROSEATE reduces the resources needed to profile temperature traits across diverse environmental samples. Beyond operational efficiencies, ROSEATE’s use of *de novo* structural embeddings ensures that predictions are not tied to knowledge about specific gene families or prior protein-specific information. This flexibility will allow the framework to be expanded to other proteins with distinct structural dynamics, even in proteins from poorly characterized bacteria. Furthermore, both by using learned representations of proteins which incorporate residue contexts and comparing the learned representations used in the OGT predictor to biophysical patterns in proteins, ROSEATE offers a more mechanistically interpretable alternative to genome-based heuristics like dipeptide frequencies. Further improvements to these structural comparisons can provide independent biophysical evidence for limitations on protein activity and stability which influence the distributions of organisms that cannot otherwise be readily studied *in vitro*. Collectively, the ROSEATE framework provides a scalable, structurally informed predictive method with particular efficiency advantages in the field of microbial ecology, offering significant advantages as phenotypic datasets and predictive methods continue to mature.

## Methods

### Compilation of protein and OGT dataset

The Gosha database [72] was used to identify bacteria with optimal growth temperatures (OGT) and associated genomes. 9,352 records from Gosha matched species representatives from the Genome Taxonomy Database (GTDB) v214.1 [73]. The 9,352 GTDB representative genomes were selected to analyze the structural properties of adenylate kinase (ADK) across temperature optima. An ADK sequence was identified in 9,303 out of 9,352 genomes using the Pfam family PF00406 HMMER model with hmmscan v3.4 [107] with a minimum bitscore of 100. The highest-scoring protein sequences (multiple hits were only found in 655 genomes and no hits were found in 49 genomes) were used to construct a dataset of 9,303 putative ADK proteins. Each protein in this dataset was folded using ColabFold v1.5.5 [82], and manually quality-filtered by length, and structural confidence (pLDDT) using custom Python and SLURM scripts. 8,532 protein structures were retained after quality filtering. The main reasons for removal were incorrectly annotated homologous proteins, misfolded proteins, and incomplete proteins. 27 structural features were computed using a combination of Rosetta v362 [108] and FreeSASA v2.2.1 [109] to assess whether canonical thermoadaptive structural features computed from the ADKs were correlated with OGT. The 8,352 proteins produced by these procedures were the proteins used to develop the ESM-based approach ROSEATE: beginning with AlphaFold structure determination ensured that each protein in our OGT prediction efforts had a quality-controlled three-dimensional structure which could be used for later validation.

### PLM embeddings

MSA Transformer [70] was used to generate flexible numeric representations of changes in ADK protein structure across taxa. An alignment for input into MSA Transformer was computed using 8,352 quality-controlled ADKs from representative GTDB genomes and 701 curated eukaryotic, archaeal, and bacterial ADKs from protein family TIGR01351 included as an outgroup. The proteins were dereplicated using CD-HIT v4.8.1 [110] with options “-c 0.95 -n 5 - M 0 -T 0” and initially aligned with MAFFT v7.525 [111] using the L-INS-i algorithm and 1000 iterations. Sequences with more than 15% gaps or large numbers of rare gaps were progressively eliminated from the alignment via manual filtering; all sequences which were present in the final filtered, dereplicated data (6,330) were re-aligned with MAFFT using L-INS-I and 1000 iterations. This finalized “backbone” alignment was trimmed with trimal v1.5.rev0 [112] in gappyout mode before all original sequences absent from the backbone (2,202) were reinserted using MAFFT with –keeplength –add. This process ensured the backbone alignment was built from a set of quality-controlled, non-redundant sequences and represented only conserved ADK residues representing each of the 8,352 ADKs with high-quality AlphaFold structures. The finalized alignment was loaded using custom Python code for embedding with the pretrained 12-layer ESM-1b MSA Transformer model (esm_msa1b_t12_100M_UR50S) installed via the ESM v2.0.0 [113] conda package was used to embed the residues of each protein in a L x 768-unit embedding vector where L = 215 for the backbone MSA; each embedding vector is a complex numeric representation which captures the identity and biophysical attributes of each residue’s unique context across the protein. All embeddings used randomly selected 8 sequences as context.

### Training and testing datasets

The OGT values for the 9,351 bacterial genomes from Gosha used to construct training and testing datasets (Fig S2A). To ensure an even sampling of OGT values between 0 and 80°C and assess the impact of dataset size on model performance, two partitions of the curated 8,352 ADK proteins were constructed. The first partition contained 1,227 unique ADK sequences evenly sampled from the range of OGT values, including some which may be of uncertain quality. This partition is hereafter referred to as dataset E, or ‘Even’ dataset (Fig S2B-C). The second partition, composed of 230 records which match the bacteria from [7] with experimentally determined growth curve data, is referred to as dataset C, or the ‘Corkrey’ dataset (Fig S2D).

### Predictive modeling using embeddings from MSA Transformer

We tested a number of models, including regression trees, linear models, Gaussian process regressors, and neural networks, either using the CLS feature (embedding component 0, representing the full protein) or per-residue embeddings to predict OGT. Each model was trained using one of the C or E partitions of the Gosha dataset and tested using the other. In both cases, the testing dataset was embedded using the training dataset’s sequences as context.

Random forests (RFs) and support vector regressions (SVRs) implemented in scikit-learn v1.4.1.post1 [114] had hyperparameters tuned with hyperopt v0.2.7 [115] used negative mean squared loss and 5-fold cross validation. Parameters tuned were n_estimators (100-500), max_depth (3-30), min_samples_split (0.01-0.2), max_features (sqrt, log2, n_features), and criterion (squared error, absolute error) for RandomForestRegressor and C (1e-3-1e3), kernel (rbf, linear, poly), gamma (1e-4-1e1), epsilon (0.01-0.5), and degree (2-5) for SVR. Weighted least-square regressions in statsmodels v0.14.1 [116] for both the CLS and per-site embeddings were fitted; a per-site model using only highly correlated sites was also considered. Based on the findings from [117], sklearn-based Gaussian process regressors were fitted using a similar CLS-or per-site-based approach using a Matern / WhiteNoise kernel and ten random restarts. Neural networks implemented in PyTorch v2.3.0 [118] used a regressor containing 3 linear layers and 2 ReLu layers with dropout = 0.1 to transform 768-unit input embeddings to 1-unit outputs. To accommodate per-site embeddings in a per-site model, an attention layer with two linear layers and a TanH activation was used to train per-site weights. The neural networks were trained using the Adam optimizer, learning rates of 1e-3 to 1e-5, and 200 epochs. The individual accuracies of these models are presented in Table S2.

Validations using variations on ROSEATE’s architecture were used to ensure that the model was not deriving predictive power from unexpected sources. A one-hot encoded model and ROSEATE models using embeddings extracted from layer 1 and layer 6 of MSA Transformer were compared to ROSEATE. These models were implemented identically to ROSEATE, using identical training and test datasets, and identical contexts for embedding. ESM likelihoods for each protein were computed by summing log soft-maxed logits across residues for each protein.

### Comparison of ROSEATE and TOME

To construct a TOME model which used the same training data as ROSEATE, TOME v1.0.0 [45] was installed from GitHub. Using TOME functions, dipeptide frequencies for the 1,227 genomes in the ‘Even’ dataset were computed, and the TOME model was retrained on this dataset. The proteomes of the organisms in the ‘Corkrey’ dataset were retrieved from GTDB, and the TOME model retrained on the ‘Even’ dataset was tested using these data. TOME retrained with ‘Even’ was also used in comparisons between ROSEATE and TOME on the datasets of novel isolate genomes described in the “Application of model to genomic and metagenomic datasets” section.

### Phylogenetic nearest-neighbors model and cross-validation procedure

A phylogenetic nearest-neighbors model was constructed to assess the influence of phylogenetic covariation on temperature optima and construct a performance baseline for further models. Using ete3 v3.1.2 [119] the GTDB phylogeny was pruned to the 1,457 genomes in the ‘Even’ and ‘Corkrey’ datasets with preserve_branch_lengths=True. The ‘Corkrey’ dataset was used as a test dataset; the OGT prediction for each test genome was the OGT of the non-test (‘Even’ dataset) genome with the shortest patristic distance.

To assess the influence of phylogenetic distances on the performance of ROSEATE TOME, and other model types, models were retrained using the best-performing parameters from initial tests in a leave-one-out cross validation procedure structured by the phylogenetic groups in the ‘Even’ dataset. For each taxonomic level (Genus to Phylum), the genomes in each unique taxon were isolated from the rest of the dataset and used as testing data. The models were retrained on the remainder of the ‘Even’ dataset, and their performance was assessed using RMSE. This procedure yielded estimates of the accuracy of each model when tested on increasing degrees of phylogenetic dissimilarity to its training data, or the phylogenetic generalizability of the model. Assessments of this phylogenetic generalizability were used to select ElasticNet as the model framework for ROSEATE and in comparisons of ROSEATE and TOME.

### Structural linear model

A structural linear model was used to assess the capacity of structural features computed from AlphaFold structures. Weighted least-squares (WLS) regression implemented in statsmodels v0.14.1 [116] was used to perform regression on 25 structural features computed with Rosetta and FreeSASA as previously discussed. After feature selection with the sklearn-based implementation of ElasticNetCV using L1 ratios between 0.5 and 0.9, weights for individual OGT records were computed using an sklearn-based scikit-learn v1.4.1.post1 [114] inverse gaussian kernel to down-weight densely sampled regions of the OGT spectrum. The finalized WLS, fitted using weighted OGT records and the subset of features which survived ElasticNet feature selection, were used to identify structural features for further comparisons with ESM embedding components.

### Comparison of AlphaFold and PLM features

To validate the association of MSA Transformer embedding components with AlphaFold-based structural features, per-residue structural features were computed for each site in the protein. Because the protein sequences in AlphaFold structures were trimmed differently than the input MSA used for embeddings, the trimmed sequences from each AlphaFold PDB file were inserted into the backbone alignment using MAFFT and standard parameters. Per-residue polar and nonpolar SASAs were calculated using FreeSASA [109] and contact densities were computed using BioPython v1.83 [120], using the alignment of protein structures to the backbone MSA to match residues. Differences in SASA between folded and unfolded states (ΔSASA) were calculated using the empirically determined SASAs [121]. Correlations and analyses between MSA Transformer embeddings and calculated structural features were conducted using custom scripts written in Python v3.10.13.

### Application of model to genomic and metagenomic datasets

To test the performance of ROSEATE on isolates, the model was applied to two new datasets of isolates not included in Gosha. The first includes 48 *Arthrobacter* strains with experimentally measured growth optima [79]. The second dataset includes 20 isolates from the genus *Mycobacterium* with measured growth curves across a temperature gradient [78]. If necessary, genomes were assembled with SPAdes v4.2.0 [122] in isolate mode and open reading frames were annotated with Prodigal v2.6.3 [123]. ADK sequences were extracted from each genome with a minimum bitscore of 100 using the PF00406 HMMer model. Only the best hit by bitscore was retained. Each ADK was inserted into the backbone protein alignment using MAFFT –add –keeplength and embedded using random selections of sequences from the training data for the model. The resulting embeddings were used as inputs for the predictive model. For each genome, predictions were compared to experimental OGT values collected using procedures described in [79] and [78] for *Arthrobacter* and *Mycobacterium*, respectively.

A similar procedure for assembling, annotating, and incorporating short read datasets into the model was used for analyses of the nine metagenomic datasets. Shotgun metagenomic datasets (415 metagenomes in total) from European freshwater lakes [91], Arctic ocean samples [96,97], hot spring metagenomes [94], marine sediments [95], cattle rumen [92] and avian gut microbiomes [93], along with soils from Australia [89] and Antarctica [79,90] were assembled using a custom SLURM pipeline the University of Colorado Research Computing Cluster. NCBI SRA or ENA ERR archives were downloaded with sratoolkit v3.3.0 (https://github.com/ncbi/sra-tools) and quality controlled with fastp v1.0.1 [124], then assembled with SPAdes in metagenome mode. After assembly, contigs were annotated with Prodigal and ADK sequences were extracted from each metagenome assembly, aligned, and embedded, with up to 1000 ADKs extracted per metagenome using a minimum bitscore of 100. Environmental temperatures were collected from metadata associated with each of the metagenomic studies.

## Acknowledgements

We would like to thank JL Weissman and members of the Fierer Lab for their willingness to engage in productive discussions about this work.

## Data availability

This study makes use of publicly available data resources. All code and data generated in this project are available in the following GitHub repository: https://github.com/realmichaelhoffert/adenylate_kinase_ogt. Please refer to Methods for additional information on the datasets and software used.

## Supporting Information

**Table S1. Description of temperature-related structural features of proteins.** Column 1 indicates whether the study was used adenylate kinase (ADK).

**Table S2. Accuracies of the six model architectures tested.** RF = Random Forest, SVR = Support Vector Regression, OLS = ordinary least-squares, ENET / ENET CV = ElasticNet-based linear regression, NN = neural network. Each model was trained on either a small (‘Corkrey’, n = 230) high-quality dataset or a larger, lower-quality dataset (‘Even’, n = 1227), see Methods for details. Best-performing models are bolded, with lower ranks indicating better performance. * = model only using highly correlated sites. RMSE = Root mean squared error, R^2^ = variance explained, and CV = phylogenetic cross-validation performance. CV rank was assigned using average error and maximum error calculated in a phylogenetic cross-validation procedure.

**Fig S1. Comparison of optimal growth temperatures with structure and structural features of adenylate kinase (ADK).** (A) Schematic diagram of the structure of adenylate kinase (ADK) generated from *E. coli* ADK (PDB: 1AKE) illustrating the highly conserved core (helices a1, a5, a6, and a7, sheets s1-5) and NMP-binding components (helices a2-4), and the partially conserved lid domain. (B-D) Relationships of ADK structural characteristics to OGT for 8,532 quality-controlled ADK structures, with Spearman R coefficients indicated as “SpR”. (B) OGT versus lid domain type for four ADK isoforms: lidded ADKs containing a 4C zinc-binding motif (Zinc), lidded ADKs containing a HSDT motif (H-bond), lidded ADKs containing neither Zinc or H-bond motifs, and lidless ADKs. Letters indicate significantly different groups in a Holm-corrected ANOVA using Welch’s t-tests with unequal variance. (C) Average total, polar, and nonpolar solvent-accessible surface areas (SASAs) and difference in SASA between folded and unfolded states (ΔSASA). (D) Additional correlated structural characteristics: the score of the ref2015 Rosetta energy function, total carbon-carbon (CC) contacts, and contact density at 4.5 angstroms.

**Fig S2. Descriptions of OGT datasets used in model training and testing.** (A) Diagram of divisions of the original 9,351 annotated ADKs through quality-filtering and division into ‘Corkrey’ (C) and ‘Even’ (E) datasets (see Results and Methods). (B) Distribution of OGT values across the Gosha, C, and E datasets, including the percent of records in Gosha with OGT equal to 20°C, 28°C, 37°C, and all other OGT values (*). (C) Phylogenetic distribution of taxa and corresponding OGT values for ‘Even’ dataset. (D) Phylogenetic distribution of taxa and corresponding OGT values for the ‘Corkrey’ dataset. (E) Comparison of OGT records for the same taxa between a robust dataset derived from growth curves (7) on the x-axis and matching records from Gosha on the y-axis showing that Gosha contains both discretized and potentially erroneous records.

**Fig S3. Comparison of the ROSEATE model to TOME using novel datasets of isolates.** (A) Accuracy of ROSEATE and TOME for a dataset of 20 strains from the genus *Mycobacterium*. (B) Accuracy of ROSEATE and TOME for a dataset of 48 strains of the genus *Arthrobacter* isolated from Antarctic (blue) and non-Antarctic (orange) soils.

**Fig S4. Results of phylogenetic cross-validation (CV) for ElasticNet, used in ROSEATE, and TOME.** Using the best-performing parameters from initial testing, an instance of each model type was trained using a leave-one-out phylogenetic cross-validation approach on the ‘Even’ dataset. For each taxon at five taxonomic levels, the model was fitted on all data except individuals from a single target taxon (leave-one-out). The model’s error (RMSE) was measured using this remaining taxon and repeated across all taxa at all levels. Red dashed line shows error of 10°C. Red dots and labels in each plot show phylum-level average CV error for each model type.

**Fig S5. Performance of additional sequence-based models for validation of ROSEATE.** (A) OGT values for versus MSA Transformer log-likelihoods for the “Even” dataset, computed as the sum of logits across each protein from the layer 12 outputs. (B) Accuracy of a “one-hot” model averaged across all sites applied to the “Corkrey” dataset. The one-hot model uses an identical architecture to ROSEATE but substitutes per-site embeddings for per-site encodings of residue identity in a one-hot matrix.

**Fig S6. Layer-specific model accuracies.** Spearman R correlations of per-site model predictions with OGT for ROSEATE and two identical models using embeddings from the first and sixth layer of MSA Transformer. As the embeddings pass through the MSA Transformer the attention modules in each layer successively adjust per-site embeddings to capture specific elements of the interactions present at each site. This consideration of interaction effects, presumably including those which influence thermostability, improve the predictive performance of models using the embeddings.

**Fig S7. Components of a generalized linear model used to quantify relationships between structural relationships and OGT.** (A) Fitted OGT values for each protein in the dataset versus actual OGT values from a generalized linear model fitted on structural features of ADK with a y = x line marked with red dashes. (B) Coefficients of the generalized linear model are colored blue for negative values and red for positive values. Abbreviations: BB = backbone, S = sidechain, SASA = solvent accessible surface area, Main = main chain, side = side chain, ΔSASA = difference in surface area between folded and unfolded states, Packing statistic = Rosetta’s packing statistic. All statistics expect SASAs were measured with Rosetta (102). (C) Residual values, with vertical gray dashed lines showing common temperatures for cultivation of bacterial isolates: 20, 25, 28, 30, 35, and 37°C, a horizontal red dashed line indicating residual values of zero, and solid gray indicating smoothed average residual value.

## References

1. Andersson S, Nilsson SI. Influence of pH and temperature on microbial activity, substrate availability of soil-solution bacteria and leaching of dissolved organic carbon in a mor humus. Soil Biol Biochem. 2001;33: 1181–1191. doi:10.1016/S0038-0717(01)00022-0

2. Papatheodorou EM, Argyropoulou MD, Stamou GP. The effects of large- and small-scale differences in soil temperature and moisture on bacterial functional diversity and the community of bacterivorous nematodes. Appl Soil Ecol. 2004;25: 37–49. doi:10.1016/S0929-1393(03)00100-8

3. Rinnan R, Rousk J, Yergeau E, Kowalchuk GA, Bååth E. Temperature adaptation of soil bacterial communities along an Antarctic climate gradient: predicting responses to climate warming. Glob Change Biol. 2009;15: 2615–2625. doi:10.1111/j.1365-2486.2009.01959.x

4. Ruan Y, Kuzyakov Y, Liu X, Zhang X, Xu Q, Guo J, et al. Elevated temperature and CO2 strongly affect the growth strategies of soil bacteria. Nat Commun. 2023;14: 391. doi:10.1038/s41467-023-36086-y

5. Pold G, Domeignoz-Horta LA, Morrison EW, Frey SD, Sistla SA, DeAngelis KM. Carbon Use Efficiency and Its Temperature Sensitivity Covary in Soil Bacteria. mBio. 2020;11: 10.1128/mbio.02293-19. doi:10.1128/mbio.02293-19

6. Alster CJ, Weller ZD, von Fischer JC. A meta-analysis of temperature sensitivity as a microbial trait. Glob Change Biol. 2018;24: 4211–4224. doi:10.1111/gcb.14342

7. Corkrey R, McMeekin TA, Bowman JP, Ratkowsky DA, Olley J, Ross T. The Biokinetic Spectrum for Temperature. PLOS ONE. 2016;11: e0153343. doi:10.1371/journal.pone.0153343

8. Dill KA, Ghosh K, Schmit JD. Physical limits of cells and proteomes. Proc Natl Acad Sci. 2011;108: 17876–17882. doi:10.1073/pnas.1114477108

9. Alster CJ, Koyama A, Johnson NG, Wallenstein MD, von Fischer JC. Temperature sensitivity of soil microbial communities: An application of macromolecular rate theory to microbial respiration. J Geophys Res Biogeosciences. 2016;121: 1420–1433. doi:10.1002/2016JG003343

10. Alster CJ, von Fischer JC, Allison SD, Treseder KK. Embracing a new paradigm for temperature sensitivity of soil microbes. Glob Change Biol. 2020;26: 3221–3229. doi:10.1111/gcb.15053

11. García FC, Clegg T, O’Neill DB, Warfield R, Pawar S, Yvon-Durocher G. The temperature dependence of microbial community respiration is amplified by changes in species interactions. Nat Microbiol. 2023;8: 272–283. doi:10.1038/s41564-022-01283-w

12. Oliverio AM, Bradford MA, Fierer N. Identifying the microbial taxa that consistently respond to soil warming across time and space. Glob Change Biol. 2017;23: 2117–2129. doi:10.1111/gcb.13557

13. Bodor A, Bounedjoum N, Vincze GE, Erdeiné Kis Á, Laczi K, Bende G, et al. Challenges of unculturable bacteria: environmental perspectives. Rev Environ Sci Biotechnol. 2020;19: 1–22. doi:10.1007/s11157-020-09522-4

14. Clarke A. Costs and consequences of evolutionary temperature adaptation. Trends Ecol Evol. 2003;18: 573–581. doi:10.1016/j.tree.2003.08.007

15. Clarke A. Is there a Universal Temperature Dependence of metabolism? Funct Ecol. 2004;18: 252–256. doi:10.1111/j.0269-8463.2004.00842.x

16. Corkrey R, Olley J, Ratkowsky D, McMeekin T, Ross T. Universality of Thermodynamic Constants Governing Biological Growth Rates. PLOS ONE. 2012;7: e32003. doi:10.1371/journal.pone.0032003

17. Venev SV, Zeldovich KB. Thermophilic Adaptation in Prokaryotes Is Constrained by Metabolic Costs of Proteostasis. Mol Biol Evol. 2018;35: 211–224. doi:10.1093/molbev/msx282

18. Vartoukian SR, Palmer RM, Wade WG. Strategies for culture of ‘unculturable’ bacteria. FEMS Microbiol Lett. 2010;309: 1–7. doi:10.1111/j.1574-6968.2010.02000.x

19. Arbab S, Ullah H, Khan MIU, Khattak MNK, Zhang J, Li K, et al. Diversity and distribution of thermophilic microorganisms and their applications in biotechnology. J Basic Microbiol. 2022;62: 95–108. doi:10.1002/jobm.202100529

20. Hector TE, Sgrò CM, Hall MD. Thermal limits in the face of infectious disease: How important are pathogens? Glob Change Biol. 2021;27: 4469–4480. doi:10.1111/gcb.15761

21. Tourte M, Schaeffer P, Grossi V, Oger PM. Membrane adaptation in the hyperthermophilic archaeon Pyrococcus furiosus relies upon a novel strategy involving glycerol monoalkyl glycerol tetraether lipids. Environ Microbiol. 2022;24: 2029–2046. doi:10.1111/1462-2920.15923

22. Ernst R, Ejsing CS, Antonny B. Homeoviscous Adaptation and the Regulation of Membrane Lipids. J Mol Biol. 2016;428: 4776–4791. doi:10.1016/j.jmb.2016.08.013

23. Maffucci I, Laage D, Stirnemann G, Sterpone F. Differences in thermal structural changes and melting between mesophilic and thermophilic dihydrofolate reductase enzymes. Phys Chem Chem Phys. 2020;22: 18361–18373. doi:10.1039/D0CP02738C

24. Zeldovich KB, Berezovsky IN, Shakhnovich EI. Protein and DNA Sequence Determinants of Thermophilic Adaptation. PLOS Comput Biol. 2007;3: e5. doi:10.1371/journal.pcbi.0030005

25. Butterwick JA, Patrick Loria J, Astrof NS, Kroenke CD, Cole R, Rance M, et al. Multiple Time Scale Backbone Dynamics of Homologous Thermophilic and Mesophilic Ribonuclease HI Enzymes. J Mol Biol. 2004;339: 855–871. doi:10.1016/j.jmb.2004.03.055

26. Somero GN. Adaptation of enzymes to temperature: searching for basic “strategies.” Comp Biochem Physiol B Biochem Mol Biol. 2004;139: 321–333. doi:10.1016/j.cbpc.2004.05.003

27. Atkinson JT, Jones AM, Nanda V, Silberg JJ. Protein tolerance to random circular permutation correlates with thermostability and local energetics of residue-residue contacts. Protein Eng Des Sel. 2019;32: 489–501. doi:10.1093/protein/gzaa012

28. Lipscomb GL, Hahn EM, Crowley AT, Adams MWW. Reverse gyrase is essential for microbial growth at 95 °C. Extremophiles. 2017;21: 603–608. doi:10.1007/s00792-017-0929-z

29. Vettone A, Perugino G, Rossi M, Valenti A, Ciaramella M. Genome stability: recent insights in the topoisomerase reverse gyrase and thermophilic DNA alkyltransferase. Extremophiles. 2014;18: 895–904. doi:10.1007/s00792-014-0662-9

30. Forterre P, Bergerat A, Lopex-Garcia P. The unique DNA topology and DNA topoisomerases of hyperthermophilic archaea. FEMS Microbiol Rev. 1996;18: 237–248. doi:10.1111/j.1574-6976.1996.tb00240.x

31. Verma D, Kumar V, Satyanarayana T. Genomic attributes of thermophilic and hyperthermophilic bacteria and archaea. World J Microbiol Biotechnol. 2022;38: 135. doi:10.1007/s11274-022-03327-z

32. Sengupta P, Garrity P. Sensing temperature. Curr Biol. 2013;23: R304–R307. doi:10.1016/j.cub.2013.03.009

33. Aguilar PS, Hernandez-Arriaga AM, Cybulski LE, Erazo AC, de Mendoza D. Molecular basis of thermosensing: a two-component signal transduction thermometer in Bacillus subtilis. EMBO J. 2001;20: 1681–1691. doi:10.1093/emboj/20.7.1681

34. Inda ME, Vandenbranden M, Fernández A, de Mendoza D, Ruysschaert J-M, Cybulski LE. A lipid-mediated conformational switch modulates the thermosensing activity of DesK. Proc Natl Acad Sci. 2014;111: 3579–3584. doi:10.1073/pnas.1317147111

35. Yamamoto M, Arai H, Ishii M, Igarashi Y. Role of two 2-oxoglutarate:ferredoxin oxidoreductases in Hydrogenobacter thermophilus under aerobic and anaerobic conditions. FEMS Microbiol Lett. 2006;263: 189–193. doi:10.1111/j.1574-6968.2006.00415.x

36. Berg IA. Ecological Aspects of the Distribution of Different Autotrophic CO2 Fixation Pathways. Appl Environ Microbiol. 2011;77: 1925–1936. doi:10.1128/AEM.02473-10

37. Knapp BD, Huang KC. The Effects of Temperature on Cellular Physiology. Annu Rev Biophys. 2022;51: 499–526. doi:10.1146/annurev-biophys-112221-074832

38. Parks DH, Chaumeil P-A, Mussig AJ, Rinke C, Chuvochina M, Hugenholtz P. GTDB release 10: a complete and systematic taxonomy for 715 230 bacterial and 17 245 archaeal genomes. Nucleic Acids Res. 2026;54: D743–D754. doi:10.1093/nar/gkaf1040

39. Sauer DB, Wang D-N. Predicting the optimal growth temperatures of prokaryotes using only genome derived features. Bioinformatics. 2019;35: 3224–3231. doi:10.1093/bioinformatics/btz059

40. Aptekmann AA, Nadra AD. Core promoter information content correlates with optimal growth temperature. Sci Rep. 2018;8: 1313. doi:10.1038/s41598-018-19495-8

41. Cimen E, Jensen SE, Buckler ES. Building a tRNA thermometer to estimate microbial adaptation to temperature. Nucleic Acids Res. 2020;48: 12004–12015. doi:10.1093/nar/gkaa1030

42. Jensen DB, Vesth TC, Hallin PF, Pedersen AG, Ussery DW. Bayesian prediction of bacterial growth temperature range based on genome sequences. BMC Genomics. 2012;13: S3. doi:10.1186/1471-2164-13-S7-S3

43. Jensen SE, Johnson LC, Casstevens T, Buckler ES. Predicting protein domain temperature adaptation across the prokaryote-eukaryote divide. bioRxiv; 2021. p. 2021.07.13.452245. doi:10.1101/2021.07.13.452245

44. Kurokawa M, Higashi K, Yoshida K, Sato T, Maruyama S, Mori H, et al. Metagenomic Thermometer. bioRxiv; 2022. p. 2022.07.14.499854. doi:10.1101/2022.07.14.499854

45. Li G, Rabe KS, Nielsen J, Engqvist MKM. Machine Learning Applied to Predicting Microorganism Growth Temperatures and Enzyme Catalytic Optima. ACS Synth Biol. 2019;8: 1411–1420. doi:10.1021/acssynbio.9b00099

46. Barnum TP, Crits-Christoph A, Molla M, Carini P, Lee HH, Ostrov N. Predicting microbial growth conditions from amino acid composition. 2024. doi:10.1101/2024.03.22.586313

47. Toki S, Matsui M, Tominaga K, Suzuki TK, Tsuchimatsu T, Iwasaki W. Genomic and evolutionary factors influencing the prediction accuracy of optimal growth temperature in prokaryotes. mSystems. 2026;11: e00062–26. doi:10.1128/msystems.00062-26

48. Colette S, François J, De Moor B, van Noort V. OGTFinder: A Curated Growth Temperature Data Set and Its Application To Predict Optimal Growth Temperatures of Bacteria and Archaea. J Chem Inf Model. 2026 [cited 24 Apr 2026]. doi:10.1021/acs.jcim.5c03033

49. Hu E-Z, Lan X-R, Liu Z-L, Gao J, Niu D-K. A positive correlation between GC content and growth temperature in prokaryotes. BMC Genomics. 2022;23: 110. doi:10.1186/s12864-022-08353-7

50. Taverna DM, Goldstein RA. Why are proteins marginally stable? Proteins Struct Funct Bioinforma. 2002;46: 105–109. doi:10.1002/prot.10016

51. Somero GN. Temperature Adaptation of Enzymes: Biological Optimization Through Structure-Function Compromises. Annu Rev Ecol Syst. 1978;9: 1–29. doi:10.1146/annurev.es.09.110178.000245

52. Fields PA, Dong Y, Meng X, Somero GN. Adaptations of protein structure and function to temperature: there is more than one way to ‘skin a cat.’ Podrabsky JE, Stillman JH, Tomanek L, editors. J Exp Biol. 2015;218: 1801–1811. doi:10.1242/jeb.114298

53. Kumar S, Tsai C-J, Nussinov R. Factors enhancing protein thermostability. Protein Eng Des Sel. 2000;13: 179–191. doi:10.1093/protein/13.3.179

54. Stirnemann G, Sterpone F. Mechanics of Protein Adaptation to High Temperatures. J Phys Chem Lett. 2017;8: 5884–5890. doi:10.1021/acs.jpclett.7b02611

55. van Noort V, Bradatsch B, Arumugam M, Amlacher S, Bange G, Creevey C, et al. Consistent mutational paths predict eukaryotic thermostability. BMC Evol Biol. 2013;13: 7. doi:10.1186/1471-2148-13-7

56. Klipcan L, Safro I, Temkin B, Safro M. Optimal growth temperature of prokaryotes correlates with class II amino acid composition. FEBS Lett. 2006;580: 1672–1676. doi:10.1016/j.febslet.2006.02.013

57. Gress A, Kalinina OV. SphereCon—a method for precise estimation of residue relative solvent accessible area from limited structural information. Bioinformatics. 2020;36: 3372–3378. doi:10.1093/bioinformatics/btaa159

58. Wang G-Z, Lercher MJ. Amino acid composition in endothermic vertebrates is biased in the same direction as in thermophilic prokaryotes. BMC Evol Biol. 2010;10: 263. doi:10.1186/1471-2148-10-263

59. Dzeja P, Terzic A. Adenylate Kinase and AMP Signaling Networks: Metabolic Monitoring, Signal Communication and Body Energy Sensing. Int J Mol Sci. 2009;10: 1729–1772. doi:10.3390/ijms10041729

60. Bae E, Phillips GN. Roles of static and dynamic domains in stability and catalysis of adenylate kinase. Proc Natl Acad Sci. 2006;103: 2132–2137. doi:10.1073/pnas.0507527103

61. Bae E, Phillips GN. Structures and Analysis of Highly Homologous Psychrophilic, Mesophilic, and Thermophilic Adenylate Kinases *. J Biol Chem. 2004;279: 28202–28208. doi:10.1074/jbc.M401865200

62. Moon S, Kim J, Bae E. Structural analyses of adenylate kinases from Antarctic and tropical fishes for understanding cold adaptation of enzymes. Sci Rep. 2017;7: 16027. doi:10.1038/s41598-017-16266-9

63. Shibanuma Y, Nemoto N, Yamamoto N, Sampei G-I, Kawai G. Crystal structure of adenylate kinase from an extremophilic archaeon Aeropyrum pernix with ATP and AMP. J Biochem (Tokyo). 2020;168: 223–229. doi:10.1093/jb/mvaa043

64. Segall-Shapiro TH, Nguyen PQ, Dos Santos ED, Subedi S, Judd J, Suh J, et al. Mesophilic and Hyperthermophilic Adenylate Kinases Differ in Their Tolerance to Random Fragmentation. J Mol Biol. 2011;406: 135–148. doi:10.1016/j.jmb.2010.11.057

65. Rao R, Meier J, Sercu T, Ovchinnikov S, Rives A. Transformer protein language models are unsupervised structure learners. bioRxiv; 2020. p. 2020.12.15.422761. doi:10.1101/2020.12.15.422761

66. Zhang Z, Wayment-Steele HK, Brixi G, Wang H, Kern D, Ovchinnikov S. Protein language models learn evolutionary statistics of interacting sequence motifs. Proc Natl Acad Sci. 2024;121: e2406285121. doi:10.1073/pnas.2406285121

67. Peng C, Bian Y, Yuan C, Chen Y, Liu D, Lu F, et al. ESM-PsyPred: Leveraging Protein Language Models for Accurate Prediction of Psychrophilic Proteins. Interdiscip Sci Comput Life Sci. 2026 [cited 21 May 2026]. doi:10.1007/s12539-025-00810-7

68. Muir DF, Grosjean P, Pinney MM, Keiser MJ. Leveraging Protein Language Model Embeddings for Catalytic Turnover Prediction of Adenylate Kinase Orthologs in a Low-Data Regime. arXiv; 2025. doi:10.48550/arXiv.2505.03066

69. Muir DF, Asper GPR, Notin P, Posner JA, Marks DS, Keiser MJ, et al. Evolutionary-scale enzymology enables exploration of a rugged catalytic landscape. Science. 2025;388: eadu1058. doi:10.1126/science.adu1058

70. Rao RM, Liu J, Verkuil R, Meier J, Canny J, Abbeel P, et al. MSA Transformer. Proceedings of the 38th International Conference on Machine Learning. PMLR; 2021. pp. 8844–8856. Available: https://proceedings.mlr.press/v139/rao21a.html

71. Jansson JK, Hofmockel KS. The soil microbiome — from metagenomics to metaphenomics. Curr Opin Microbiol. 2018;43: 162–168. doi:10.1016/j.mib.2018.01.013

72. Helena-Bueno K, Brown CR, Melnikov S. Gosha: a database of organisms with defined optimal growth temperatures. 2021. doi:10.1101/2021.12.21.473645

73. Parks DH, Chuvochina M, Rinke C, Mussig AJ, Chaumeil P-A, Hugenholtz P. GTDB: an ongoing census of bacterial and archaeal diversity through a phylogenetically consistent, rank normalized and complete genome-based taxonomy. Nucleic Acids Res. 2022;50: D785–D794. doi:10.1093/nar/gkab776

74. Detlefsen NS, Hauberg S, Boomsma W. Learning meaningful representations of protein sequences. Nat Commun. 2022;13: 1914. doi:10.1038/s41467-022-29443-w

75. Schmirler R, Heinzinger M, Rost B. Fine-tuning protein language models boosts predictions across diverse tasks. Nat Commun. 2024;15: 7407. doi:10.1038/s41467-024-51844-2

76. Eom H, Park S, Cho KS, Lee J, Kim H, Kim S, et al. Discovery of highly active kynureninases for cancer immunotherapy through protein language model. Nucleic Acids Res. 2025;53: gkae1245. doi:10.1093/nar/gkae1245

77. Peng C, Bian Y, Yuan C, Chen Y, Liu D, Lu F, et al. ESM-PsyPred: Leveraging Protein Language Models for Accurate Prediction of Psychrophilic Proteins. Interdiscip Sci Comput Life Sci. 2026 [cited 24 Apr 2026]. doi:10.1007/s12539-025-00810-7

78. Gebert MJ, Lipner EM, Galletta JM, Henley JB, Hoffert M, Riskin ML, et al. A cross-environment comparison of nontuberculous mycobacterial diversity. Appl Environ Microbiol. 2025;92: e01552–25. doi:10.1128/aem.01552-25

79. Dragone NB, Childress MK, Mendez N, Galletta J, Vanderburgh C, Mesquita CPB de, et al. Evidence for endemism and local adaptation in Antarctic soil bacteria. bioRxiv; 2026. p. 2026.05.01.722257. doi:10.64898/2026.05.01.722257

80. Roberts DR, Bahn V, Ciuti S, Boyce MS, Elith J, Guillera-Arroita G, et al. Cross-validation strategies for data with temporal, spatial, hierarchical, or phylogenetic structure. Ecography. 2017;40: 913–929. doi:10.1111/ecog.02881

81. Albright S, Louca S. Trait biases in microbial reference genomes. Sci Data. 2023;10: 84. doi:10.1038/s41597-023-01994-7

82. Mirdita M, Schütze K, Moriwaki Y, Heo L, Ovchinnikov S, Steinegger M. ColabFold: making protein folding accessible to all. Nat Methods. 2022;19: 679–682. doi:10.1038/s41592-022-01488-1

83. Jiang F, Li M, Dong J, Yu Y, Sun X, Wu B, et al. A general temperature-guided language model to design proteins of enhanced stability and activity. Sci Adv. 2024;10: eadr2641. doi:10.1126/sciadv.adr2641

84. Vig J, Madani A, Varshney LR, Xiong C, Socher R, Rajani NF. BERTology Meets Biology: Interpreting Attention in Protein Language Models. arXiv; 2021. doi:10.48550/arXiv.2006.15222

85. Gilles AM, Glaser P, Perrier V, Meier A, Longin R, Sebald M, et al. Zinc, a structural component of adenylate kinases from gram-positive bacteria. J Bacteriol. 1994;176: 520–523. doi:10.1128/jb.176.2.520-523.1994

86. Saavedra HG, Wrabl JO, Anderson JA, Li J, Hilser VJ. Dynamic allostery can drive cold adaptation in enzymes. Nature. 2018;558: 324–328. doi:10.1038/s41586-018-0183-2

87. Moon S, Kim J, Koo J, Bae E. Structural and mutational analyses of psychrophilic and mesophilic adenylate kinases highlight the role of hydrophobic interactions in protein thermal stability. Struct Dyn. 2019;6: 024702. doi:10.1063/1.5089707

88. Alford RF, Leaver-Fay A, Jeliazkov JR, O’Meara MJ, DiMaio FP, Park H, et al. The Rosetta All-Atom Energy Function for Macromolecular Modeling and Design. J Chem Theory Comput. 2017;13: 3031–3048. doi:10.1021/acs.jctc.7b00125

89. Bissett A, Fitzgerald A, Meintjes T, Mele PM, Reith F, Dennis PG, et al. Introducing BASE: the Biomes of Australian Soil Environments soil microbial diversity database. GigaScience. 2016;5: 21. doi:10.1186/s13742-016-0126-5

90. Dragone NB, Childress MK, Vanderburgh C, Willmore R, Hogg ID, Sancho LG, et al. A comprehensive survey of soil microbial diversity across the Antarctic continent. Polar Biol. 2025;48: 50. doi:10.1007/s00300-025-03372-y

91. Shah M, Bornemann TLV, Nuy JK, Hahn MW, Probst AJ, Beisser D, et al. Genome-resolved metagenomics reveals the effect of nutrient availability on bacterial genomic properties across 44 European freshwater lakes. Environ Microbiol. 2024;26: e16634. doi:10.1111/1462-2920.16634

92. Conteville LC, da Silva JV, Andrade BGN, Cardoso TF, Bruscadin JJ, de Oliveira PSN, et al. Rumen and fecal microbiomes are related to diet and production traits in Bos indicus beef cattle. Front Microbiol. 2023;14: 1282851. doi:10.3389/fmicb.2023.1282851

93. Youngblut ND, de la Cuesta-Zuluaga J, Reischer GH, Dauser S, Schuster N, Walzer C, et al. Large-Scale Metagenome Assembly Reveals Novel Animal-Associated Microbial Genomes, Biosynthetic Gene Clusters, and Other Genetic Diversity. mSystems. 2020;5: e01045–20. doi:10.1128/mSystems.01045-20

94. Colman DR, Keller LM, Arteaga-Pozo E, Andrade-Barahona E, St. Clair B, Shoemaker A, et al. Covariation of hot spring geochemistry with microbial genomic diversity, function, and evolution. Nat Commun. 2024;15: 7506. doi:10.1038/s41467-024-51841-5

95. Mara P, Geller-McGrath D, Edgcomb V, Beaudoin D, Morono Y, Teske A. Metagenomic profiles of archaea and bacteria within thermal and geochemical gradients of the Guaymas Basin deep subsurface. Nat Commun. 2023;14: 7768. doi:10.1038/s41467-023-43296-x

96. Royo-Llonch M, Sánchez P, Ruiz-González C, Salazar G, Pedrós-Alió C, Sebastián M, et al. Compendium of 530 metagenome-assembled bacterial and archaeal genomes from the polar Arctic Ocean. Nat Microbiol. 2021;6: 1561–1574. doi:10.1038/s41564-021-00979-9

97. Salazar G, Paoli L, Alberti A, Huerta-Cepas J, Ruscheweyh H-J, Cuenca M, et al. Gene Expression Changes and Community Turnover Differentially Shape the Global Ocean Metatranscriptome. Cell. 2019;179: 1068–1083.e21. doi:10.1016/j.cell.2019.10.014

98. van Gestel NC, Ducklow HW, Bååth E. Comparing temperature sensitivity of bacterial growth in Antarctic marine water and soil. Glob Change Biol. 2020;26: 2280–2291. doi:10.1111/gcb.15020

99. Cao Z, Zhang H, Zou Z. Language models reveal a complex sequence basis for adaptive convergent evolution of protein functions. Proc Natl Acad Sci. 2025;122: e2418254122. doi:10.1073/pnas.2418254122

100. Zhu M, Song Y, Yuan Q, Yang Y. Accurately predicting optimal conditions for microorganism proteins through geometric graph learning and language model. Commun Biol. 2024;7: 1709. doi:10.1038/s42003-024-07436-3

101. Jiang F, Li M, Dong J, Yu Y, Sun X, Wu B, et al. A general temperature-guided language model to design proteins of enhanced stability and activity. Sci Adv. 2024;10: eadr2641. doi:10.1126/sciadv.adr2641

102. Engqvist MKM. Correlating enzyme annotations with a large set of microbial growth temperatures reveals metabolic adaptations to growth at diverse temperatures. BMC Microbiol. 2018;18: 177. doi:10.1186/s12866-018-1320-7

103. Madin JS, Nielsen DA, Brbic M, Corkrey R, Danko D, Edwards K, et al. A synthesis of bacterial and archaeal phenotypic trait data. Sci Data. 2020;7: 170. doi:10.1038/s41597-020-0497-4

104. Sato Y, Okano K, Kimura H, Honda K. TEMPURA: Database of Growth TEMPeratures of Usual and RAre Prokaryotes. Microbes Environ. 2020;35: ME20074. doi:10.1264/jsme2.ME20074

105. Bains W, Xiao Y, Yu C. Prediction of the Maximum Temperature for Life Based on the Stability of Metabolites to Decomposition in Water. Life. 2015;5: 1054–1100. doi:10.3390/life5021054

106. Jaenicke R. Stability and stabilization of globular proteins in solution. J Biotechnol. 2000;79: 193–203. doi:10.1016/s0168-1656(00)00236-4

107. Finn RD, Clements J, Eddy SR. HMMER web server: interactive sequence similarity searching. Nucleic Acids Res. 2011;39: W29–W37. doi:10.1093/nar/gkr367

108. Leman JK, Weitzner BD, Lewis SM, Adolf-Bryfogle J, Alam N, Alford RF, et al. Macromolecular modeling and design in Rosetta: recent methods and frameworks. Nat Methods. 2020;17: 665–680. doi:10.1038/s41592-020-0848-2

109. Mitternacht S. FreeSASA: An open source C library for solvent accessible surface area calculations. F1000Research; 2016. doi:10.12688/f1000research.7931.1

110. Fu L, Niu B, Zhu Z, Wu S, Li W. CD-HIT: accelerated for clustering the next-generation sequencing data. [cited 19 Feb 2026]. Available: 10.1093/bioinformatics/bts565

111. Katoh K, Standley DM. MAFFT Multiple Sequence Alignment Software Version 7: Improvements in Performance and Usability. Mol Biol Evol. 2013;30: 772–780. doi:10.1093/molbev/mst010

112. Capella-Gutiérrez S, Silla-Martínez JM, Gabaldón T. trimAl: a tool for automated alignment trimming in large-scale phylogenetic analyses. [cited 19 Mar 2025]. Available: 10.1093/bioinformatics/btp348

113. Lin Z, Akin H, Rao R, Hie B, Zhu Z, Lu W, et al. Evolutionary-scale prediction of atomic-level protein structure with a language model. Science. 2023;379: 1123–1130. doi:10.1126/science.ade2574

114. Pedregosa F, Pedregosa F, Varoquaux G, Varoquaux G, Org N, Gramfort A, et al. Scikit-learn: Machine Learning in Python. Mach Learn PYTHON.

115. Bergstra J, Komer B, Eliasmith C, Yamins D, Cox DD. Hyperopt: a Python library for model selection and hyperparameter optimization. Comput Sci Discov. 2015;8: 014008. doi:10.1088/1749-4699/8/1/014008

116. Seabold S, Perktold J. Statsmodels: Econometric and Statistical Modeling with Python. Austin, Texas; 2010. pp. 92–96. doi:10.25080/Majora-92bf1922-011

117. Michael R, Kæstel-Hansen J, Groth PM, Bartels S, Salomon J, Tian P, et al. A systematic analysis of regression models for protein engineering. PLOS Comput Biol. 2024;20: e1012061. doi:10.1371/journal.pcbi.1012061

118. Paszke A, Gross S, Massa F, Lerer A, Bradbury J, Chanan G, et al. PyTorch: An Imperative Style, High-Performance Deep Learning Library. Advances in Neural Information Processing Systems. Curran Associates, Inc.; 2019. Available: https://proceedings.neurips.cc/paper/2019/hash/bdbca288fee7f92f2bfa9f7012727740-Abstract.html

119. Huerta-Cepas J, Serra F, Bork P. ETE 3: Reconstruction, Analysis, and Visualization of Phylogenomic Data. Mol Biol Evol. 2016;33: 1635–1638. doi:10.1093/molbev/msw046

120. Cock PJA, Antao T, Chang JT, Chapman BA, Cox CJ, Dalke A, et al. Biopython: freely available Python tools for computational molecular biology and bioinformatics. Bioinformatics. 2009;25: 1422–1423. doi:10.1093/bioinformatics/btp163

121. Tien MZ, Meyer AG, Sydykova DK, Spielman SJ, Wilke CO. Maximum Allowed Solvent Accessibilites of Residues in Proteins. PLOS ONE. 2013;8: e80635. doi:10.1371/journal.pone.0080635

122. Prjibelski A, Antipov D, Meleshko D, Lapidus A, Korobeynikov A. Using SPAdes De Novo Assembler. Curr Protoc Bioinforma. 2020;70: e102. doi:10.1002/cpbi.102

123. Hyatt D, Chen G-L, LoCascio PF, Land ML, Larimer FW, Hauser LJ. Prodigal: prokaryotic gene recognition and translation initiation site identification. BMC Bioinformatics. 2010;11: 119. doi:10.1186/1471-2105-11-119

124. Chen S. fastp 1.0: An ultra-fast all-round tool for FASTQ data quality control and preprocessing. [cited 2 Mar 2026]. doi:10.1002/imt2.70078

